# Evolution of the mutational process under relaxed selection in *Caenorhabditis elegans*

**DOI:** 10.1101/280826

**Authors:** Ayush Shekhar Saxena, Matthew P. Salomon, Chikako Matsuba, Shu-Dan Yeh, Charles F. Baer

**Affiliations:** Department of Biology, University of Florida, Gainesville, FL, USA; Department of Molecular Oncology, John Wayne Cancer Institute, Santa Monica, CA, USA; Department of Life Sciences, National Central University, Taiwan; University of Florida Genetics Institute

## Abstract

The mutational process varies at many levels, from within genomes to among taxa. Many mechanisms have been linked to variation in mutation, but understanding of the evolution of the mutational process is rudimentary. Physiological condition is often implicated as a source of variation in microbial mutation rate and may contribute to mutation rate variation in multicellular organisms.

Deleterious mutations are a ubiquitous source of variation in condition. We test the hypothesis that the mutational process depends on the underlying mutation load in two groups of *Caenorhabditis elegans* mutation accumulation (MA) lines that differ in their starting mutation loads. “First-Order MA” (O1MA) lines maintained under minimal selection for ∼250 generations were divided into high-fitness and low-fitness groups and sets of “second-order MA” (O2MA) lines derived from each O1MA line were maintained for ∼150 additional generations. Genomes of 48 O2MA lines and their progenitors were sequenced. There is significant variation among O2MA lines in base-substitution rate (*µ*_*bs*_), but no effect of initial fitness, whereas the indel rate is greater in high-fitness O2MA lines. Overall, *µ*_*bs*_ is positively correlated with recombination and proximity to short tandem repeats and negatively correlated with 10 bp and 1 Kb GC content. However, probability of mutation is well-predicted by the three-nucleotide motif. ∼90% of the variance in standing nucleotide variation is explained by mutability. Total mutation rate increased in the O2MA lines, as predicted by the “drift barrier” model of mutation rate evolution. These data, combined with experimental estimates of fitness, suggest that epistasis is synergistic.

## Introduction

The evolution of the mutation rate is of longstanding interest to evolutionary theorists (Fisher 1930; Sturtevant 1937; Lynch et al. 2016), and there is abundant empirical evidence that the overall rate, molecular spectrum, and phenotypic consequences of mutation - collectively, the mutational process - vary at many biological levels, from within an individual genome to among species and higher taxa (Drake et al. 1998; Conrad et al. 2011; Schrider et al. 2013; Long et al. 2016; Ness et al. 2016; Carlson et al. 2018). The mechanistic, environmental, ecological, and evolutionary factors that potentially influence variation in the mutational process are legion.

Microbiologists have appreciated for many decades that physiological stress is often associated with increased mutation rate (e.g., see Figure 6 of Ogur et al. 1960). Recently, Agrawal and his colleagues have undertaken a systematic investigation into the effects of physiological condition (∼ “stress”) on the mutational process in *Drosophila melanogaster*, motivated by theoretical findings that if the mutation rate is condition-dependent, the accumulation of deleterious mutations can have interesting and sometimes counterintuitive feedback effects on population mean fitness (Agrawal 2002; Shaw and Baer 2011). They manipulated physiological condition both exogenously, by manipulating food quality (Agrawal and Wang 2008) and endogenously, by allowing mutations to accumulate under relaxed selection on genomes that were initially identical except for the presence or absence of one or two mutations of large deleterious effect (Sharp and Agrawal 2012). Poorly-fed females transmitted ∼30% more lethal mutations than did well-fed females (Agrawal and Wang 2008). Similarly, mutation accumulation (MA) lines beginning with a large genetic load declined in fitness more rapidly than lines with wild-type genomes, which is most simply explained by the low-fitness lines having a greater mutation rate than the high-fitness lines (Sharp and Agrawal 2012). Whole-genome sequencing revealed that the faster decline in fitness of the loaded MA lines can be attributed to an elevated rate of small deletions, plausibly as a result of different mechanisms of double-strand break repair (Sharp and Agrawal 2016). In an analogous study, Ávila et al. (2006) constructed a set of MA lines of *D. melanogaster* derived from a single MA line that had itself accumulated mutations under relaxed selection for 265 generations, a protocol that we call “second-order MA” (O2MA). The per-generation decline in fitness was greater in the second-order MA lines than in the ancestral (“first-order MA”, O1MA) lines. That result is consistent with an increased mutation rate in the O2MA lines relative to the O1MA lines, but it is also consistent with mutational effects being greater in the second-order MA lines (i.e., synergistic epistasis; Dickinson 2008).

We report here the results of a second-order MA experiment in the nematode *C. elegans*, specifically designed to assess the relationship between the initial genomic load of spontaneous deleterious mutations and the subsequent effects on the mutational process (Figure 1). Initially, a set of 100 MA lines derived from a single, highly inbred individual of the N2 strain was allowed to accumulate mutations for approximately 250 generations under minimal selection. From a subset of 67 O1MA lines assayed for absolute fitness (defined as lifetime reproductive success under non-competitive conditions), we chose five lines with consistently high fitness and five lines with consistently low fitness, from which we established ten independent sets of 48 O2MA lines, referred to as “O2MA families”, which were allowed to accumulate mutations for an additional ∼150 generations (Matsuba et al. 2012). Upon completion of the second-order MA phase, five replicate O2MA lines from each of the ten O2MA families were sequenced at ∼25X average coverage, along with nine of the ten O1MA progenitors and an additional 23 O1MA lines not included in the O2MA experiment.

The resulting data allow us to address several fundamental questions about the evolution of the mutational process. First, does initial fitness affect the mutational process, and if so, how? Second, how fast does genetic variation in the mutational process accumulate due to the effects of new spontaneous mutations? Third, is the mutation rate itself subject to a mutational bias, as predicted by the “drift barrier” hypothesis of mutation rate evolution? Fourth, how does the mutational process depend on underlying features of the genome, e.g., local sequence context, recombination rate or base composition? Fifth, to what extent does the local mutational milieu predict standing nucleotide sequence variation?

Finally, in combination with experimental estimates of fitness, we can assess how the average mutational effect depends on the underlying genetic load. If mutational effects are greater in the low-fitness O2MA lines than in the high-fitness lines, it implies that deleterious mutations interact synergistically; the converse implies diminishing-returns epistasis.

## Results

Pooled over 48 (out of 50) O2MA lines and nine of the ten O2MA progenitors (three lines yielded poor sequencing coverage), using our default pipeline we identified 1828 base substitutions (1481 O2MA, 347 O1MA), 361 small deletions (293 O2 MA, 68 O1MA) and 236 small insertions (196 O2MA, 40 O1MA). We sequenced the genomes of an additional 23 O1MA lines not included in the O2MA experiment, from which we identified 884 base substitutions, 142 deletions, and 93 insertions. Mutation rates for the different groups are summarized in Table 1, and given for individual lines in Supplementary Table S1. Individual mutations and their properties are listed in Supplementary Table S2. Raw sequence data are archived in the NCBI Short Read Archive, project numbers PRJNA395568 (O2MA lines and O2MA progenitors) and PRJNA429972 (other O1MA lines). Averaged over all 32 O1MA lines, the per-nucleotide base-substitution mutation rate *µ*_*bs*_ = 2.33 (± 0.08) x 10^-9^ per generation. *µ*_*bs*_ for the nine O2MA progenitors is 2.26 ± 0.12 x 10^-9^ per generation). Averaged over the 48 O2MA lines, the base-substitution mutation rate over the subsequent ∼150 generations is estimated to be *µ*_*bs*_ = 2.57 (± 0.11) x10^-9^ per-generation, not significantly different from the combined O1MA rate (general linear model, F_1,14.5_ = 1.32, p>0.26; see Supplementary Appendix A1.6 for details of the GLM). The short indel rate for the full set of 32 O1MA lines *µ*_*INDEL*_ =0.66 (±0.04) x 10^-9^ per-site per-generation. The indel rate of the nine O2MA progenitors (*µ*_*INDEL,ANC*_ = 0.68 ± 0.06 x 10^-9^/generation) does not differ from that of the other 21 O1MA lines (*µ*_*INDEL,OTHER*_ = 0.64 ± 0.05 x 10^-9^/generation). Averaged over all 48 O2MA lines, *µ*_*INDEL*_ = 0.84 ± 0.05 x10^-9^/generation, significantly greater than the combined O1MA rate (GLM, F_1,24.2_ = 8.27, p < 0.01) but not significantly greater than that of the nine O2MA progenitors (GLM, F_1,45.4_ = 1.61, p > 0.21).

**Figure 1.**
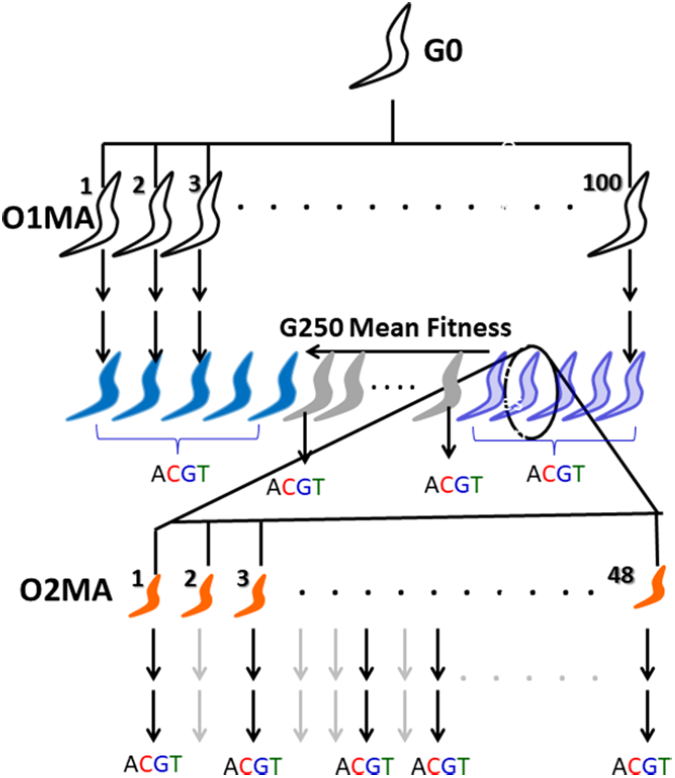
Schematic depiction of the second-order MA (O2MA) experiment. After ∼250 generations of MA (unfilled worms), five low-fitness O1MA lines (light blue worms) and five high-fitness O1MA lines (dark blue worms) were chosen as progenitors (“O2MA progenitor”, circled worm) for sets of 48 O2MA lines (orange worms). Each set of 48 O2MA lines derived from an O2MA progenitor is an “O2MA family”. After ∼150 additional generations (G400), five O2MA lines were randomly chosen from each O2MA family for sequencing (“ACGT”). At the same time, the O2MA progenitor of each O2MA family was sequenced, along with 23 randomly chosen O1MA lines that were not progenitors of O2MA families (gray worms).

**Table 1.** Average mutation rates (SEM).

|  | O2MA<br>progenitor (*n*=9) | O1MA<br>other (*n*=23) | O1MA<br>all (*n*=32) | O2MA, High<br>(*n*=24) | O2MA, Low<br>(*n*=24) | O2MA, all<br>(*n*=48) |
| --- | --- | --- | --- | --- | --- | --- |
| $\mu_{bs} (\times 10^9)$ | 2.26 (0.12) | 2.35 (0.11) | 2.33 (0.08) | 2.58 (0.11) | 2.49 (0.32) | 2.57 (0.11) |
| $\mu_{INS} (\times 10^9)$ | 0.26 (0.04) | 0.25 (0.03) | 0.26<sup>x</sup> (0.02) | 0.35 (0.03) | 0.32 (0.04) | 0.34<sup>y</sup> (0.03) |
| $\mu_{DEL} (\times 10^9)$ | 0.42 (0.05) | 0.39 (0.04) | 0.40 (0.03) | 0.59<sup>a</sup> (0.07) | 0.41<sup>b</sup> (0.04) | 0.50 (0.04) |
| $\mu_{INDEL} (\times 10^9)$ | 0.68 (0.06) | 0.64 (0.05) | 0.66<sup>x</sup> (0.04) | 0.95<sup>a</sup> (0.06) | 0.73<sup>b</sup> (0.05) | 0.84<sup>y</sup> (0.05) |
| $\mu_{Total} (\times 10^9)$ | 2.94 (0.11) | 2.99 (0.13) | 2.98<sup>x,*</sup> (0.10) | 3.52 (0.13) | 3.21 (0.31) | 3.37<sup>y,*</sup> (0.16) |
| $\mu_{GENOME}$ | 0.29 | 0.30 | 0.30 | 0.35 | 0.32 | 0.34 |

All mutation rates are per-site, per-generation except $\mu_{GENOME}$ . Abbreviations are: $\mu_{bs}$ , base substitution mutation rate; $\mu_{INS}$ , insertion rate; $\mu_{DEL}$ , deletion rate; $\mu_{INDEL}$ , indel rate; $\mu_{Total}$ , total mutation rate; $\mu_{GENOME}$ , haploid genome-wide mutation rate per-generation. <sup>a,b</sup> superscripts are significantly different (*P*<0.05) within O1MA or O2MA groups. <sup>x,y</sup> superscripts are significantly different (*P*<0.05) between O1MA and O2MA. \* *P*≈0.06.

There is a positive correlation between the number of callable sites and the estimated base-substitution mutation rate *µ*_*bs*_ (*r*_*HIGH,O2MA*_ = 0.25, p < 0.22, *r*_*LOW,O2MA*_ = 0.58, p < 0.002, *r*_*O1MA*_ = 0.30, p < 0.09). In the O2MA lines, callable sites are confounded with both O2MA family and fitness. However, the correlation is driven by a few lines with atypically low coverage (average number of callable sites less than 60% of the genome). The indel rate is uncorrelated with callable sites (*r*_*HIGH,O2MA*_ = 0.16, p > 0.44, *r*_*LOW,O2MA*_ = 0.12, p > 0.58, *r*_*O1MA*_ = 0.00, p > 0.99). We elaborate on potential artifactual and/or biological factors that may contribute to the correlation in the Extended Discussion in Supplementary Appendix A2.1. We do not believe the results are meaningfully affected by the correlation between *µ*_*bs*_ and callable sites.

### No evidence for fitness-dependent base-substitution mutation

There is significantly more variation in *µ*_*bs*_ among O2MA lines than expected if the base-substitution mutation rate is uniform across the full set of 48 O2MA lines (simulation P<0.0001; Supplementary Figure S1; see Methods and Supplementary Appendix 3.1 for details of simulations). Moreover, there is significant variation in *µ*_*bs*_ among O2MA families (LRT, chi-square = 9.27, df=1, P<0.003; Figure 2). Similarly, there is more variation among O1MA lines than predicted by a uniform mutation rate (simulation P<0.015). The simplest interpretation is that some element(s) of the mutational process diverged over the course of the first ∼250 generations of MA, and that the signal of the difference(s) carried through the next ∼150 generations of O2MA.

However, there is no evidence that *µ*_*bs*_ differs consistently between the O2MA families derived from high fitness and low fitness O2MA progenitors (GLM, F_1,6.24_ = 0.18, p>0.75;). Averaged over the five O2MA families in each fitness class, the base substitution mutation rate between G250 and G400 for High and Low fitness lines is estimated to be *µ_bs,HIGH_* = 2.58 (±0.11) x 10^-9^/ gen, and *µ_bs,LOW_* = 2.49 (±0.32) x 10^-9^ / gen (Figure 2).

As expected from the lack of differentiation of *µ*_*bs*_, the base-substitution spectrum does not differ significantly between the high-fitness and low-fitness O2MA lines (Supplementary Figure S2; Monte Carlo Fisher’s Exact Test, 10^7^ replicates, P>0.70), nor does it vary between O2MA families (MC FET, P>0.25), between individual O2MA lines (MC FET, P>0.08), between O2MA lines within any of the ten families (P>0.10 or greater in all ten cases) or between the O2MA progenitors at G250 and the O2MA lines at G400 (Supplementary Figure S2; MC FET, P>0.51).

Consistent with many previous studies (Lynch 2007), the average mutation rate from a C or G to an A or T is significantly greater than the mutation rate from A or T to C or G (*µ*_*C/G*→*A/T*_ = 3.03 (±0.18) x 10^-9^/gen; *µ*_*A/T*→*C/G*_ = 0.93 (±0.05) x 10^-9^/gen). Extrapolating from these rates, the expected base composition of the *C. elegans* genome at mutational equilibrium is ∼76.5% AT, greater than the actual AT fraction of ∼64.5%. Inspection of the *µ*_*bs*_ data reveals two potential outlying O2MA lines (Figure 2), one low fitness (O2MA line 508.34, *µ*_*bs*_ = 0.78 x 10^-9^/gen) and one high fitness (O2MA line 579.36, *µ*_*bs*_ = 4.34 x 10^-9^/gen). When those two lines are omitted from the analysis, the variance among O2MA lines is sufficiently explained by a single base-substitution rate (simulation P>0.07).

However, O2MA lines derived from O2MA progenitor 508 have the lowest base-substitution rate even with the extreme line omitted [*µ*_*bs*_ = 1.49 (±1.90) x 10^-9^/gen with line 508.34 included, 1.68 (±0.91) x 10^-9^/gen) without line 508.34] and O2MA lines derived from O2MA progenitor 579 have the highest base-substitution rate even with the extreme line omitted [*µ*_*bs*_ = 3.53 (±0.31) x 10^-9^/gen with line 579.36 included, 3.26 (± 0.22) x 10^-9^/gen) without line 579.36]. The random chance that the most extreme high line comes from the family whose other four members also have the highest average mutation rate (5/48) and that the most extreme low line comes from the family whose other four members also have the lowest average mutation rate (5/47) is approximately 1%. The most parsimonious explanation is that the two outlying O2MA lines are simply the most extreme manifestations of a biological process common to their respective O2MA progenitors rather than true outliers. The alternative is that the apparently extreme mutation rates are experimental artifacts, which we think is unlikely (see Extended Discussion in Supplementary Appendix A2).

**Figure 2.**
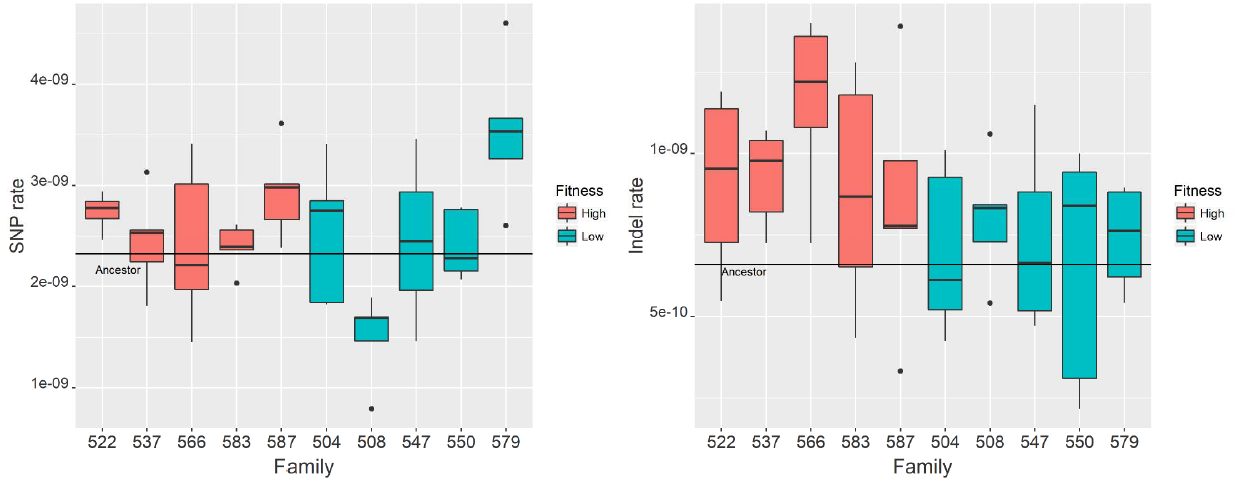
Distribution of mutation rate among O2MA families. Families derived from high-fitness O2MA progenitors are in orange, families derived from low-fitness O2MA progenitors are in green. Families 522 and 547 contain four sequenced O2MA lines, the other families contain five O2MA lines. The horizontal line denotes the mutation rate of the nine O2MA progenitors. Points shown outside the box are beyond the 1.5X inter-quartile range of the family whereas whiskers represent data points within that range. that the mutational process in the ten O2MA progenitors diverged by ∼250 generations of evolution under MA conditions, and the signal of the difference(s) was retained over the subsequent ∼150 generations of O2MA. (A, left) Base-substitution mutation rate (*µ*_*bs*_). (B, right) Indel mutation rate (*µ*_*INDEL*_).

### Fitness-dependence of the small indel rate

In contrast to the base-substitution mutation rate, which does not differ between O2MA lines derived from high fitness and low fitness O1MA lines, the O2MA short indel rate is significantly greater in the high fitness group (*µ*_*INDEL*_ = 0.95 ± 0.06 x10^-9^/generation) than in the low fitness group (*µ*_*INDEL*_ = 0.73 ± 0.50 x10^-^/generation: GLM, F_1,44.4_ = 7.84, P<0.01). The difference is primarily due to different rates of deletions (*µ*_*DEL,High*_ = 0.59 ± 0.07 x10^-9^/generation, *µ_DEL,Low_* = 0.41 ± 0.04 x10^-9^/generation; F_1,7.58_ = 5.77, P < 0.05) rather than insertions (*µ*_*INS,High*_ = 0.35 ± 0.03 x10^-9^/generation, *µ_INS,Low_* = 0.32 ± 0.04 x10^-9^/generation; F_1,17.5_ = 0.51, P > 0.48). The variance in *µ*_*INDEL*_ among O2MA lines within each fitness group is adequately explained by a single, fitness-specific indel rate (high-fitness, simulation P>0.12; low-fitness, simulation P>0.2). The distribution of indel lengths is given in Supplementary Figure S3.

The higher indel rate of high-fitness O2MA lines suggests that the indel rate of O1MA lines should be greater than the overall O2MA rate, given the higher fitness of the G0 ancestor. Interestingly, that is not what we observe. The overall O1MA indel rate, including all 32 O1MA lines, is significantly lower than high-fitness O2MA indel rate (*µ*_*INDEL, O1MA*_ = 0.66 ± 0.04 x10-^9^/generation, *µ*_*INDEL, O2MA_High*_ = 0.95 ± 0.06 x10^-10^/generation; GLM, F_1,41.7_=15.61, P<0.0005), but not significantly different from low fitness O2MA indel rate (*µ*_*INDEL, O1MA*_ = 0.66 ± 0.04 x10-^9^/generation, *µ*_*INDEL, O2MA_Low*_ = 0.73 ± 0.05 x10^-9^/generation; F_1,47.6_ = 1.10, P>0.29). The results do not change if only the nine O2MA progenitors are used to calculate the indel rate.

The O2MA insertion and deletion rates are uncorrelated (*r_INS,DEL_* = 0.021, P>0.88, *n*=48), suggestive of different factors underlying the two types of mutations. The base-substitution rate is moderately positively correlated with the insertion rate (*r_bs,INS_* = 0.32, P<0.03, *n*=48) but uncorrelated with the deletion rate (*r_bs,DEL_* = −0.016, P>0.91, *n*=48).

There are two potential evolutionary factors that can explain the difference in the deletion rate between the different O2MA fitness groups. First, some element of the mutational process may differ, e.g., DNA repair. Alternatively, selection may differ in either strength or efficiency between the two treatments. In our MA protocol, differences in selection efficiency between lines can result from different frequencies of going to backup (see Methods). However, the mean effectively neutral selection coefficient (*s*_*n*_ =1/4*N_e_*) is only slightly smaller in the low-fitness lines 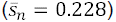 than in the high-fitness lines 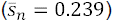. The slightly lower efficiency of selection in the high fitness group seems unlikely to account for the ∼30% greater indel rate.

Alternatively, the strength of selection itself may be different between the two fitness groups, such that some mutations that are only mildly deleterious – and thus effectively neutral – in a high-fitness line are significantly more deleterious in a low-fitness line, i.e., epistasis is synergistic (= negative; Phillips 2008). In that case, a larger fraction of mutations in the low-fitness lines would exceed the threshold of effective neutrality and be removed by selection. The mutational effect analysis, as implemented in snpEff 4.1 (see Methods), reveals no significant difference in the frequency of potentially large-effect indels (20/303 among high-fitness O2MA lines, 10/186 among low-fitness lines, Fisher’s Exact Test, two-tailed P>0.70) or SNPs (22/822 vs. 15/659, Yates’ Chi-square, two-tailed P>0.75). Nor does the frequency of potentially large-effect mutations differ between the high and low-fitness G250 O1MA lines, between high-fitness O2MA and O1MA lines, or between low-fitness O2MA and O1MA lines (P>0.12 in all cases). The preceding analysis provides no information about mutations that did not fix. Nevertheless, synergistic epistasis, coupled with a slight overall increase in mutation rate, seems a more parsimonious explanation than different indel rates in high-fitness and low-fitness O2MA lines. Experimentally derived estimates of relative fitness (Matsuba et al. 2012) combined with estimates of the average number of mutations carried by high-fitness and low-fitness O2MA lines reinforce this conclusion (see Discussion).

### Mutation is positively associated with recombination

Standing nucleotide diversity is almost always positively associated with recombination rate, which could in principle be due either to the mutagenicity of recombination or to natural selection (i.e., Hill-Robertson interference). There is good reason to think that HR interference has an important causal role in this pattern (Lynch 2007). However, with the important exception of humans, most of the data are derived from inferences drawn from comparisons with a reference class of genomic sites believed to be free from selective constraints (e.g., processed pseudogenes or four-fold degenerate sites) rather than directly from *de novo* mutations, and the extent to which recombination is mutagenic *per se* remains an open question. Several large studies of *de novo* mutations in humans report a significant association between recombination rate and mutation rate (Michaelson et al. 2012; Francioli et al. 2015; Carlson et al. 2018), although recombination appears to affect nucleotide diversity beyond its association with mutation. Two cohort studies in bees (Yang et al. 2015; Liu et al. 2017) found a weak relationship between recombination and mutation, but the numbers of mutations were small and the inferences somewhat circumstantial. Conversely, Ness et al. (2015) found no association between recombination rate and mutation rate in the green alga *Chlamydomonas reinhardtii*, although the data on recombination rate in *C. reinhardtii* are sparser than for humans. In *C. elegans*, the relative proportions of different types of variants (SNPs, small and large indels) in the standing genetic variation differ depending on the local recombination rate, which has been suggested to reflect the signature of recombination-dependent mutation (Hwang and Wang 2017).

To investigate the relationship between recombination and mutation, we determined the association between *µ*_*bs*_ estimated from the O2MA lines and recombination rate using weighted OLS regression, dividing each chromosome into the recombination rate bins reported by Rockman and Kruglyak (2009) and weighting each bin by its size in Mb. Recombination rate is nearly constant within each bin (see Figure 1 in Rockman and Kruglyak (2009)). Contrary to our previous report from a different (and smaller) subset of G250 O1MA lines (Denver et al. 2009), here we find a significant positive univariate association between recombination rate and *µ*_*bs*_ (pseudo-*r*^2^ = 0.26, P<0.003, Supplementary Figure S4); presumably the discrepancy between this study and the previous report is due to the greater power afforded by the much greater number of mutations included in the present study (316 vs. 1828).

### The genomic correlates of mutability

Many features of the genome and epigenome influence the mutational process. The effects of some such features are well-understood and seem to be relatively general to all living organisms (e.g., short tandem repeats, G:C vs. A:T, 5’-methyl-C), whereas others remain uncertain and/or appear to be taxon-specific. To more fully characterize the features of the *C. elegans* genome that are associated with the mutational process, we employed a logistic regression method, loosely following the approach of Michaelson et al. (2012) and Ness et al. (2015). Since the deletion rate differs significantly between the two O2MA fitness groups, we restricted the analysis to base-substitutions. Univariate logistic regression coefficients of the features included in the full multiple regression are shown in Figure 3.

To assess the predictive power of the mutability model, we randomly sampled half of the O2MA lines to train the model (24 lines; roughly 740 mutations) and tested the model on the remaining 24 O2MA lines. All mutant sites and 100,000 randomly chosen non-mutant sites were arbitrarily binned into 35 bins of uniform width, and the observed mutation rate for each mutability bin was plotted against expected mutability. This analysis was repeated 100 times. Of the factors initially included in the multiple regression (see Methods), the best model includes only the 64 three-base motifs as a set of predictor variables (Figure 4a). Any combination of other predictors, with or without the three-base motif, results in a poorer fit. The poorer fit presumably results from either overfitting, multicollinearity and/or the inability of the logistic regression model to accommodate non-linear relationships between predictor variables and mutation rate, even when tuned with high penalty (λ) to drop predictor variables altogether (Lasso). Logistic regression coefficients are tabulated in Supplementary Table S3.

It is reassuring but not surprising that the model provides a good fit to the MA data from which it was trained. Of more interest is the relationship between mutability as predicted from MA data and the genetic variation observed in nature. We obtained publicly-available whole genome sequence data from 40 *C. elegans* wild isolates (Thompson et al. 2013) and identified SNPs using the same pipeline that we used to call putative mutations in the MA data. We identified ∼537,000 SNPs by these criteria. Sites were categorized as variable or not variable, without regard to allele frequency.

Mutability was assessed as described previously, except in this analysis the model was trained on the full set of 1828 mutations. Figure 4b shows a plot of nucleotide diversity (quantified as θ_*W*_; (Watterson 1975)) at non-coding sites (288,585 intron sites, and 122,272 intergenic sites) against predicted mutability. Averaged over all bins, mutability strongly predicts standing nucleotide diversity, although the variance is high at predicted high-mutability sites, presumably because the sample sizes are small.

### Short Tandem Repeats (STRs)

Short tandem repeat loci (“microsatellites”) can mutate orders of magnitude faster than other classes of loci, and potentially contribute a large fraction of the per-generation mutational variance. We previously estimated the haploid per-genome mutation rate of dinucleotide STR loci in the full set of O1MA lines to be ∼0.12/generation (Phillips et al. 2009). That calculation accounts for variation in mutation rate among repeat motifs (e.g., AT vs. AG, etc.) and at least partly accounts for variation in mutation rate with repeat number, although there is substantial uncertainty that cannot be easily quantified. Seyfert et al. (2008) found no significant effect of repeat length (di, tri, or tetranucleotide repeat) on the rate of STR mutation in a different set of N2-strain MA lines. Denver et al. (2004) investigated the mutational process of mononucleotide repeats (= “homopolymers”) in the same MA lines as Seyfert et al. and concluded that the (haploid) per-genome mononucleotide mutation rate is about 0.8/generation, more than twice the rate of all other classes of mutations combined. Mutational properties of STRs are summarized by repeat type in Supplementary Table S4 and Supplementary Figures S5 and S6.

**Figure 3.**
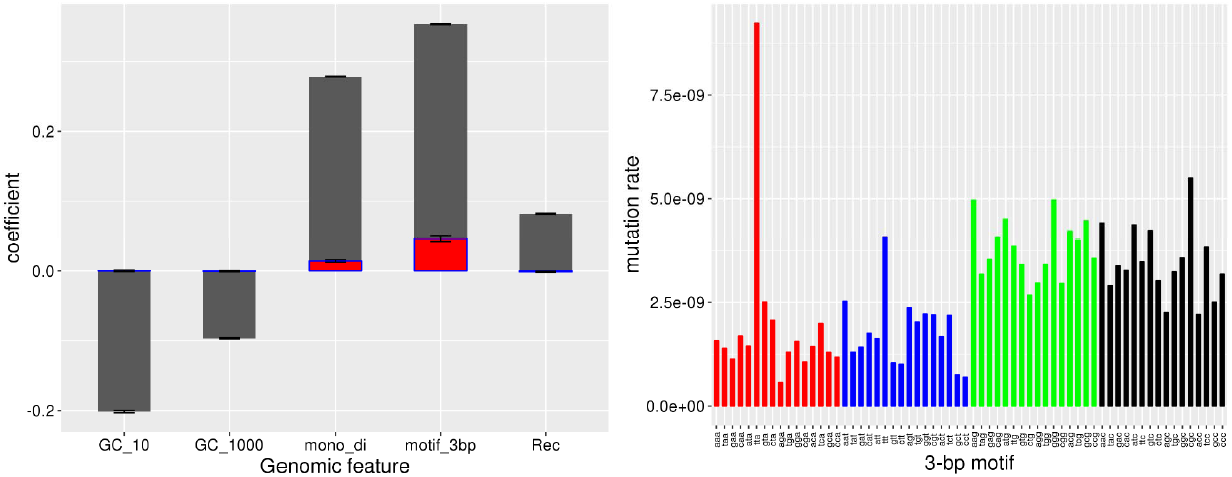
(A, Left panel) Univariate logistic regression coefficients for different genomic features (gray bars). Overlapping red bars depict the null expectation from sampling variance with 1828 randomly selected genomic sites treated as mutations. Variable abbreviations are: GC_10, 10 bp GC-content centered on the focal base; GC_1000, 1 kb GC content; mono_di, proximity to a mono or dinucleotide STR; motif_3bp, three-base motif with the mutant base at the 3’ end; Rec, local recombination rate. The method used to condense several predictors into one, for 3-base motif and mono-di STR, is described in the Methods. (B, Right panel) Base-substitution mutation rate (µ_bs_) of each 3-base motif, with mutations on the 3’ end. Motifs are grouped by the mutant base.

**Figure 4.**
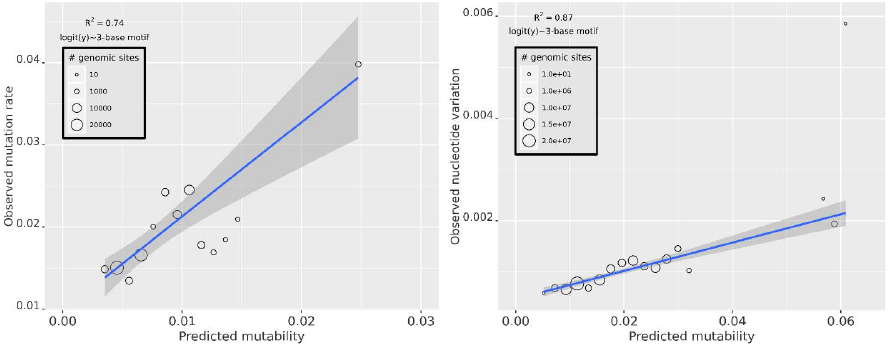
Left panel (a) depicts the relationship between observed and predicted mutability across mutability bins (n=35 bins). The mutability model includes only the 3-base motif. Right panel (b) depicts the relationship between observed standing nucleotide variation, measured as Watterson’s θ, and predicted mutability. Predicted mutability of wild isolates is calculated from the full complement of mutations in the MA lines (n=1828).

Our findings qualitatively recapitulate those of the previous studies. First, G:C mononucleotides experience indel mutations at a tenfold greater rate than A:T mononucleotides, as observed by Denver et al. (2004). Second, indel mutation rate differs only about twofold between the three A:T containing dinucleotides (AC, AG, AT) and we detected no mutations at CG dinucleotides, both as observed by Phillips et al. (2009). The ratio of mononucleotide deletions (251) to insertions (139) is not significantly different from the 16:14 ratio reported by Denver et al. (Yates’ chi-square = 1.02, df = 1, P > 0.31), and the ratio of dinucleotide deletions (10) to insertions (32) is nearly identical to the 8:28 ratio observed by Phillips et al. However, there are quantitative differences between the findings of this study and the previously reported values. The mononucleotide indel rate is estimated to be 0.056 mutations per haploid genome per generation, about 10% the rate estimated by Denver et al. (2004). The dinucleotide indel rate is estimated to be 0.003 mutations per haploid genome per generation, about 5% of the estimate from Phillips et al. (2009).

We believe the differences between the STR mutation rates estimated in this study and those reported previously are more apparent than real, and that the higher genome-wide estimates from the previous studies are probably closer to the truth. The previous studies included only a small number of STR loci, in which extremely long repeats were greatly overrepresented. In contrast, long repeats are significantly underrepresented in our data. There is no reason to doubt that mutation rate increases with repeat number. Probably, we have missed a handful of highly mutable loci that contribute a disproportionately large fraction of mutations to the genome-wide total. However, very long mono‐ and dinucleotide repeats represent a miniscule fraction of the genome (e.g., mono‐ and dinucleotides > 20 repeats represent approximately 0.02% of the genome), and it seems unlikely (although not impossible) that those few highly mutable loci contribute meaningfully to fitness. Thus, although the true genome-wide mutation rate may in fact approach one per haploid genome per generation, the meaningful mutation rate may be much lower.

We elaborate on potential causes of the source(s) of the discrepancies between this study and the previous studies in the Extended Discussion, Supplementary Appendix A2.3.

### Copy Number Variants (CNVs)

CNVs were called using the read-depth based method implemented in the CNV-seq software (Xie and Tammi 2009). The number of putative CNVs inferred is sensitive to the parameters of the analysis, with different sets of input parameters leading to estimates of the CNV rate that differ by an order of magnitude. Sensible interpretation of the results requires understanding the details of the analysis, which we outline in the Methods and elaborate in section 4 of the Extended Discussion, Appendix A2. There are two salient general results: (1) in no case does the mean number of putative CNVs differ between the high and low-fitness O2MA lines, but also (2) the number of putative CNVs in O2MA lines is consistently greater than in O1MA lines (Supplementary Table S5). Given the uncertainty associated with the estimates, we did not attempt to confirm CNVs (e.g., with qPCR). We report the results as a cautionary note that, even with as “easy” a genome as *C. elegans* N2 strain MA lines, which are almost completely homozygous and have an exceptionally well-characterized reference genome, estimates of CNVs from short-read data are extremely sensitive to methodological details. We expect that the CNV problem will eventually be resolved with accurate long-read sequencing at high coverage (Chaisson et al. 2015; Tyson et al. 2018).

## Discussion

### The Effects of Fitness. (i) Mutation Rate

There is no evidence that the rate or molecular spectrum of base-substitution mutations in the N2 strain of *C. elegans* depends on the fitness of the starting genotype. In contrast, the short indel rate, especially the deletion rate, is fitness dependent, but not in the anticipated way: low-fitness genotypes have a significantly lower deletion rate than do high-fitness genotypes. This result differs from the finding in *Drosophila melanogaster* that low-fitness genotypes experience significantly greater rates of small deletions than do high-fitness genotypes, apparently because flies in poor physiological condition employ a different, more error-prone mechanism of double-strand break repair than do flies in good condition (Sharp and Agrawal 2016).

### The Effects of Fitness. (ii) Mutational Effects

In the abstract, existence of a robust system seems to imply redundancy of components, which in the context of a genetic system implies that epistasis must be synergistic on average (de Visser et al. 2003). However, empirical evidence concerning the average epistatic effect of spontaneous deleterious mutations has been inconclusive (Halligan and Keightley 2009), and whether deleterious mutations interact synergistically *on average* has vexed generations of evolutionary biologists.

It is certainly possible that worms with high-fitness genotypes incur more short deletions than do worms with low-fitness genotypes and/or are worse at repairing them. If we use the point estimates of the average number of mutations carried by O2MA lines derived from high-fitness and low-fitness O2MA progenitors and the average mutational decline in relative fitness, averaged over all O2MA lines (Supplementary Table S6), the average mutational effect on relative fitness of the low-fitness O2MA lines is about twice that of the high fitness lines (0.48% vs 0.25%). If we assume that only indels affect fitness, the mean effects on low-fitness and high-fitness O2MA lines are about 2% and 1%, respectively. We further suppose that the “dark matter” represented by the (assumed) missing fraction of indels in the low-fitness lines would amplify the difference in average selective effects. These results represent some of the first direct evidence that spontaneous mutations interact synergistically, on average (see also Jasmin and Lenormand 2016).

### Variation in the Mutational Process

The per-generation input of genetic variance for a trait, the mutational variance, *V*_*M*_, ultimately governs the evolvability of the trait and is a fundamental parameter in evolutionary genetics (Lynch and Hill 1986). *V*_*M*_ can be estimated from MA data from the relationship 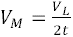, where *V*_*L*_ is the among-line component of variance and *t* is the number of generations of MA (Lynch and Walsh 1998). *V*_*M*_ is commonly standardized relative to the residual variance *V*_*E*_, called the mutational heritability 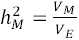. Among O2MA families, *V*_*L*_ for *µ*_*bs*_ = 0.196, V_E_ = 0.316, and 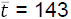, so 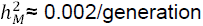. To put that result in context, 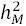 for a wide variety of traits in disparate taxa averages about 0.001/generation (Houle et al. 1996), although 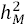 for some traits (notably gene expression) is consistently an order of magnitude less (e.g., Rifkin et al. 2005; Landry et al. 2007). That the mutation rate is evolvable is not surprising (we know it is), but the point estimate of 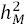 suggests that the mutational target for mutation rate is substantial. However, an estimate of a ratio of variances based on ten data points cannot be considered robust.

### Mutability and Genetic Variation

The observed positive univariate association between local recombination rate and mutation rate in this experiment is almost surely not due to Hill-Robertson interference. The expected time to fixation/loss of a new mutation is three generations (Keightley and Caballero 1997), and the average mutation rate (CNVs notwithstanding) is about one per genome every three generations. Thus, the opportunity for H-R interference in our experiment is very low, albeit not nonexistent. However, the causal factors underlying the relationship between local recombination rate and base-substitution mutation rate remain uncertain. The cause is not as simple as GC content (G:C being more mutable than A:T), because chromosome arms, which have higher recombination rates, are AT-rich. The sufficiency of 3-bp motif as a predictor of mutability suggests that the positive association of mutation with recombination is an epiphenomenon resulting from the overrepresentation of highly mutable motifs in regions of high recombination. Most clearly, mononucleotide runs are both mutagenic and positively associated with local recombination rate.

The mutability model does a good job of predicting standing nucleotide variation in natural isolates of *C. elegans*: sites that are more mutable are, on average, more variable (Figure 4b). Again, this is reassuring, but not surprising; the same relationship has been observed in *Chlamydomonas reinhardtii* (Ness et al. 2016) and humans (Francioli et al. 2015). One difference between *C. elegans* and many other organisms (e.g., humans) is that, in *C. elegans*, regions of low recombination (chromosome cores) are gene-rich rather than gene-poor, so H-R interference is more important in regions of low recombination both for inherent reasons and because the opportunity for selection is greater due to the larger target for deleterious mutations. One possibility is that, because chromosome arms are gene-poor, mutagenic features (e.g., specific motifs, such as mononucleotide runs) preferentially accumulate because their background effects on linked loci are less important. Perhaps paradoxically, the AT-richness of chromosome arms may be a signature of increased mutation rate, because C:G mutates to A:T more often than the reverse.

### Transition/Transversion ratio in MA and wild isolates

The ratio of transitions to transversions does not differ significantly between high fitness O2MA and low fitness O2MA lines (Ts/Tv_High_=0.72, Ts/Tv_Low_=0.68, *t*-test, P>0.5), nor does it differ between O2MA lines and O2MA progenitors (Ts/Tv_O1MA_=0.70, Ts/Tv_O2MA_=0.74, *t*-test, P>0.8). As in previous studies with *C. elegans* MA lines, the Ts/Tv ratio in MA lines is less than the Ts/Tv ratio in wild isolates (Denver et al. 2009; Denver et al. 2012). The difference could be due to purifying selection against transversions (Babbitt and Cotter 2011), or it could be that the mutational milieu in the lab environment differs from that in the wild. Ts/Tv of rare variants (<5% variant allele frequency relative to the reference genome, Ts/Tv = 1.20) is similar to that of common variants (>50% variant allele frequency, Ts/Tv = 1.14). That finding is not consistent with purifying selection against transversions, which suggests that some feature of the lab environment may cause the mutational spectrum of MA lines to differ from that in nature. An independent test of this possibility is to compare Ts/Tv between our N2-strain MA ancestor and the N2 reference genome; any variants will have arisen in the lab environment. In that comparison, Ts/Tv = 0.78, consistent with the MA Ts/Tv ratio and different from that among wild isolates. Denver et al. (2012) found a similarly lower Ts/Tv in MA lines of the congener *C. briggsae* relative to wild isolates, suggesting that the phenomenon is not restricted to *C. elegans*.

### Evolution of the mutation rate

The total mutation rate, *µ*_*Total*_, of the O2MA lines is about 13% greater than that of the O1MA lines (Table 1). Directional change in a trait under MA conditions (“mutational bias”, ΔM) suggests that the trait is under ongoing directional selection in the opposite direction, analogous to the direction of phenotypic change upon inbreeding (Teotónio et al. 2017). This finding is consistent with the “drift barrier” hypothesis of mutation rate evolution, which posits that directional selection to reduce the mutation rate is opposed by a weak mutational bias (Lynch 2008). It is not consistent with the mutation rate being at an optimum established by a “cost of fidelity”, wherein direct selection to reduce the input of deleterious mutations is counterbalanced by indirect selection to reduce the fitness cost of genome surveillance (Kimura 1967). If the mutation rate is at an optimum imposed by countervailing components of selection, the overall fitness function will be stabilizing. Provided that the fitness function is approximately symmetrical around the optimum, the expectation is that the among-line variance in the trait will increase but that the overall trait mean will not change. We emphasize that these findings do not imply that there is no cost of fidelity, just that the mutation rate in *C. elegans* does not appear to be at a global optimum.

The ∼13% increase in mutation rate over the course of ∼250 generations amounts to a per-generation change ΔM≈0.0005, compared to a per-generation decrease in competitive fitness of ≈0.001 in the same lines (Yeh et al. 2018). It is difficult to believe that cumulative selection on competitive fitness is only twice as strong as cumulative selection to decrease mutation rate. The most logical conclusion is that mutations that increase mutation rate have deleterious pleiotropic effects on fitness. However, this conclusion seems at odds with the failure to observe a main effect of fitness in the O2MA lines. In principle, the discrepancy could be resolved by determining the mutational correlation of mutation rate with fitness. The sample sizes necessary to answer that question in multicellular organisms are currently prohibitive, but it may be practical in a microbial system.

#### Conclusions

Sets of second-order MA lines that differed significantly in relative fitness were allowed to accumulate mutations for ∼150 generations, to test the hypothesis that physiological stress imposed by mutation load causes the mutation rate to increase. Contrary to our hypothesis, the base substitution rate did not differ significantly between high-fitness and low-fitness lines, whereas the indel rate was greater in the high-fitness lines. Averaged over all lines, the total mutation rate between generations 250-400 was significantly greater than the mutation rate between generations 0 and 250, as predicted by the “drift barrier” hypothesis of mutation rate evolution. The average mutational effect on fitness (α) is approximately twice as great in the low-fitness lines, which implies that the epistatic effects of deleterious mutations are synergistic.

### Methods and Materials

#### Mutation Accumulation Protocol

A schematic depiction of the experimental design is presented in Figure 1. Details of the first-order MA protocol and fitness assays are reported in Baer et al. (2005); details of the second-order MA protocol are reported in Matsuba et al. (2012) and summarized in the Extended Methods, Supplementary Appendix A1.1.

#### Genome sequencing and estimation of mutation rates

Five O2MA lines from the ten O2MA families and 35 O1MA lines, including the ten O2MA progenitors, were sequenced at an average of ∼25X coverage depth. Sequencing was done using Illumina technology with 100 bp paired-end reads. Protocols for DNA extraction and construction of sequencing libraries are given in Supplementary Appendix A1.2; details of preliminary processing of raw sequence data are given in Supplementary Appendix A1.3. The quality of sequence from two O2MA lines and one O1MA line was poor and these samples were omitted from further analyses, leaving 48 O2MA line and nine of the ten O2MA progenitors.

Variants were called using GATK software (McKenna et al. 2010) with a minimum coverage threshold of >10X. Variants were identified as putative mutations if (1) the variant was identified as homozygous, and (2) it was present in one and only one O2MA line. Criterion (1) means that any mutations that occurred in the last few generations of second-order MA that were still segregating and/or occurred during population expansion for DNA extraction were ignored. Because the O1MA progenitor was at mutation-drift equilibrium (Lynch and Hill 1986), the segregating variation is expected to be the same in the O1MA progenitor and the O2MA line, so ignoring heterozygotes results in an unbiased estimate of mutation rate. Criterion (2) reduces the probability of mistakenly identifying a variant segregating at low frequency in the expanded population of the O1MA progenitor as a new mutation. Two pairs of O1MA lines shared multiple variants and were inferred to have experienced contamination at some point during the MA phase; one line from each pair was arbitrarily omitted from subsequent analyses (see Extended Discussion, Supplementary Appendix A2.1).

The mutation rate (per-site, per-generation) μ of each O2MA line was calculated as *m/nt* where *m* is the number of mutations, *n* is the number of nucleotide sites observed and *t* is the number of generations of MA (Denver et al. 2009). The average mutation rate of each O1MA progenitor was calculated as the unweighted mean of the O2MA lines in that family.

We applied three additional variant-calling strategies, one with a more stringent set of criteria which we refer to as the “trimmed genome” and two with more liberal criteria, which we refer to as the “lenient genome” and “STR-relaxed”, respectively. We also explored several alternative GATK filters. Details of the alternative GATK filters are given in Supplementary Appendix A1.3; justification and details of the additional variant-calling strategies are given in Supplementary Appendix A1.4.

#### Mutation Confirmation

From each of the 48 O2MA lines, we randomly chose one putative base-substitution and one putative indel mutation for confirmation by Sanger sequencing. Details of the confirmation protocol are given in Supplementary Appendix A1.5. We confirmed 43/48 putative base-substitutions (zero false positives, five failures) and 36/48 putative indels (zero false positives, 12 failures), consistent with a false positive rate below 2.5% based on the upper 95% Poisson confidence limit. The number of failures did not differ significantly between base-substitutions and indels (Fisher’s exact test, two-tailed P>0.1).

#### Data Analysis

*i) Variation among O1MA and O2MA lines* - The simplest hypothesis regarding the mutational process is that it remained constant over the course of the experiment subsequent to the divergence of the O1MA lines from the common G0 ancestor. To test the hypothesis that a uniform mutation rate sufficiently explains variation among MA lines, we simulated the evolutionary process of 48 O2MA lines with a uniform base-substitution mutation rate equal to the unweighted mean base-substitution mutation rate of the 48 O2MA lines, accounting for the number of callable sites (Supplementary Table S2) and the number of generations of second-order MA of each line (Supplementary Table S1). At each generation, the simulated genome (100.25 Mb) of a line was assigned a number of mutations drawn from a Poisson distribution with parameter equal to the unweighted mean mutation rate. Next, each simulated line was assigned a number of generations of MA and a number of callable sites sampled with replacement from the observed distribution. Sets of 48 simulated lines were generated 100,000 times, and the observed variance in mutation rate among the 48 O2MA lines was compared to the distribution of simulated variances (Supplementary Figure S1). An analogous simulation was done for the 34 O1MA lines. Details of the simulation and code are provided in Supplementary Appendix A3.1.

To test for effects of fitness on mutation rate and to partition the variance in mutation rate into within and among-group components, we consider the mutation rate itself as a continuously-distributed dependent variable in a general linear model (GLM). Details of the GLM analyses are given in Supplementary Appendix A1.6.

It is possible that the distribution of mutation types – the mutational spectrum - differs among groups even if the overall mutation rate does not (Long et al. 2016), or if none of the type-specific mutation rates achieves statistical significance. Spectra were compared among groups by Fisher’s Exact Test using Monte Carlo sampling as implemented in the FREQ procedure of SAS v. 9.4.

*ii) Mutability* – Many factors potentially influence the probability that a site will mutate. Some of these factors can be unambiguously characterized from pooled genomic DNA from a population of multicellular organisms (local sequence motif, base composition at various scales, local recombination rate), whereas other factors that potentially influence heritable mutation are only relevant in the context of the germline or its embryonic precursors (e.g., chromatin state, nucleosome occupancy, expression level). To elucidate the relationship between the various genomic properties of a given site and the probability that a mutation occurs at that site, we employed a logistic regression model in which the log odds-ratio that a mutation occurs at a given site (success=mutation) is modeled as the sum of a set of linear predictor input variables (Michaelson et al. 2012; Ness et al. 2016). We initially included as independent variables the three-base motif with the focal nucleotide at the 3’ end and at the 5’ end (these need not be redundant if the probability that a sequencing read is included is not identical for the two strands), the five-base motif with the focal nucleotide in the center, local recombination rate (see Results for details), presence or absence of the focal site in the vicinity of a (mono/di)nucleotide run (+ 2 bp), and the 11-base and 1001-base GC-content, centered on the focal site. For cross-validation, we trained our models using half of our O2MA lines (n=24), including all base substitutions and 100,000 randomly chosen non-mutant sites, and tested the predictions on the remaining 24 O2MA lines. Following model selection, we trained the model using all the ancestral O1MA and O2MA mutations (n=1828). Including only SNPs from the O2MA lines (n=1481) had no qualitative effect on the results.

Logistic regression was performed using the GLMnet package in R (Friedman et al. 2010). The two model parameters are the tuning penalty λ and the ridge/lasso penalty α. As α→0 (ridge), the model tends to shrink the coefficients of correlated predictor variables toward each other without dropping any of the predictor variables. As α→1 (lasso), when predictor variables are correlated the model chooses one and discards the other(s). The tuning parameter λ controls the overall strength of the penalty. For all the models we tested, λ was chosen by the package’s built-in cross validation function (“lambda-min”). The fit of our models remain largely unchanged by the selection of α, with the exception of instances where α→0 (ridge). For values of α sufficiently close to 0, we observed twice the slope (bias, or regression coefficient; expected value = 1) expected when the observed mutation rate is regressed against the predicted mutation rate. All results presented here used α=0.05.

Models including short tandem repeats (STRs), five base motif and/or 10bp GC content together with three-base motif fit poorly. The poor fit could be due either to overfitting, multicollinearity and/or a non-linear relationship between a predictor variable and mutability. Moreover, models that included STRs as the only predictors fit reasonably well in terms of calibration of the model, but suffer from poorer predictive discrimination, as they lack any discrimination in the non-STR region of the genome (∼97% of the genome). The final set of models tested include only the three-base motif with the focal nucleotide at the 3’ end, 1Kb GC content and local recombination rate. For each set of input predictor variables and α, the dataset was resampled 200 times, with a different randomly chosen set of 100,000 non-mutant sites, and divided into halves, the training set and the test set. The training set consisted of half the O2MA lines, and the test set consisted of the other half. For each data partition, the model was trained and the fit to the test set was assessed by the fraction of the variance explained by the linear regression of the observed (test) value on the predicted (training) value (R^2^), the regression coefficient (bias, expected value = 1), and the area under the ROC curve (AUC, or cstatistic) for predictive discrimination (varies between 0 and 1). For each set of α and input variables, we generated a distribution of 200 R^2^ values, bias values and AUC values, and retained model(s) that maximized R^2^, and AUC while kept the regression coefficient to as close to one as possible. Code and example files are provided in Supplementary Appendix A3.2. We list the performance of all the models we tested in Supplementary Table S7.

GLMnet fits the following regression model utilizing every site in the training set, and resulting mutability is the probability of mutation “Pr (mutant)” in the following equation.

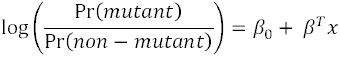

A mutability value of (say) 0.08 for a specific set of features implies that 92 out of 100 sites with the same features in the training set are non-mutant. The absolute value of mutability varies positively with the number of mutations in the training set and negatively with the number of non-mutant sites included during training. Mutability can only be interpreted in the relative context of the study. For example, Ness et al. (2015) included ∼6500 mutations in a training set with 100,000 non-mutant sites, so the average mutability in that study would be ∼3.5X greater than the estimate in this study.

Although mutability potentially varies between 0 and 1, the highest mutability observed rarely exceeded 0.2, as average mutability is the fraction of mutant sites as compared to the total number of sites. During cross-validation, we trained the model on slightly under half of the total mutations (∼740 mutants; half of all O2MA mutants, and none of the O1MA mutants), and 100,000 randomly selected sites, yielding the expected value of mutability to be ∼0.007 (740/(100000+740)). Following model selection, to assess the relationship between standing genetic variation and mutability, we recalculated mutability using all 1828 mutations. Expected mutability calculated from the full data set is ∼0.017 (1828/(100000+1828)).

To assess the relative contribution to mutability of different genomic features, we took a slightly different approach. It is not possible to condense 64 3-base motifs into a single predictor in a standard logistic regression framework. Instead, we trained a logistic regression model using only the 3-base motif, and used the predictions made from this model as predictors in a second logistic regression, which yields a single coefficient for 3-base motif. We then selected 1828 random sites from the genome, treated them as mutations, and repeated the process 100 times to obtain the null distribution for these coefficients.

*iii) Copy Number Variants* - We called putative copy-number variants (CNVs) using a read-depth (RD) based approach as implemented in the CNV-Seq software (Xie and Tammi 2009). Read-depth outliers in an MA line (O1MA or O2MA) are identified relative to its progenitor. We employed a sliding window approach, in which read-depth in a given window in the focal sample is compared to read-depth in the reference (progenitor), by taking the ratio of total reads falling in that window, normalized by the average read-depth for that chromosome in that sample. We used two different signal thresholds (1.5x/2x) for calling duplications or deletions, and several sets of sliding window sizes (see Supplementary Appendix 2, section 4 for details). The minimum detectable size of a CNV is 1kb in all cases. A putative CNV is called only when all sliding windows covering an interval meet the signal threshold. The smaller the window, the more conservative the test, because more consecutive windows must meet the signal criterion. Similarly, the greater the signal threshold, the more conservative the criterion. As with other types of variants, only putative CNVs unique to a single O2MA line are considered potential mutations. For reasons which we elaborate in the Results, we did not attempt to validate CNVs.

*iv) Predicted Mutational Effects* –To broadly categorize the distribution of predicted mutational effects, we used the publicly available software snpEff 4.1 (Cingolani et al. 2012) to annotate variants with respect to a set of characteristics potentially related to fitness (http://snpeff.sourceforge.net/VCFannotationformat_v1.0.pdf). The snpEff software uses gene lists (gtf/gff3) from the WS234 build of the *C. elegans* genome to assign a putative effect score of high, moderate, or low based on these characteristics, where a “high” effect variant is most likely to have a deleterious effect on fitness. Each variant can have multiple annotations due to different alleles and/or different splice variants; we included only the largest potential effect of each variant. Effects were parsed using custom AWK scripts and categorized as having low, medium, or high maximum impact.

## Acknowledgments

We thank Tim Crombie, Joanna Dembek, Joanna Joyner-Matos, Gigi Ostrow, Asher Shoucair and the many undergraduate students who worked over the years to maintain and propagate the MA lines. We thank Rob Ness and Jonathan Sebat for graciously explaining some details of the analyses in their papers, and Mike Miyamoto, Henrique Teotónio, and the anonymous reviewers for their helpful comments. This work was supported by NSF grant DEB-0717167 to CFB and NIH grants R01GM072639 to CFB and D. R. Denver and R01GM107227 to CFB and E. C. Andersen.

